# C-low threshold mechanoreceptor activation becomes sufficient to trigger affective pain in spinal cord-injured mice in association with increased respiratory rates

**DOI:** 10.1101/2022.11.06.515254

**Authors:** Donald J. Noble, Rochinelle Dongmo, Shangrila Parvin, Karmarcha K. Martin, Sandra M. Garraway

**Affiliations:** Department of Cell Biology, Emory University School of Medicine, Atlanta, Georgia 30322

**Author notes:** **Author Contribution Statement:** SMG and DJN designed research. DJN, SMG, SP, KKM, and RD performed research and analyzed data. DJN, SMG, SP, and KKM wrote the paper. **Correspondence Addressed To:** Sandra M. Garraway, Department of Cell Biology, Emory University School of Medicine, 615 Michael Street, Suite 605G, Atlanta, Georgia 30322. E-mail address, Donald J. Noble, Department of Cell Biology, Emory University School of Medicine, 615 Michael Street, Suite 605S, Atlanta, Georgia 30322.

**Keywords:** C-low threshold mechanoreceptors (C-LTMRs), spinal cord injury (SCI), chronic affective pain, respiratory rate (RR), Conditioned place aversion (CPA)

## Abstract

The mechanisms of neuropathic pain after spinal cord injury (SCI) are not fully understood. In addition to the plasticity that occurs within the injured spinal cord, peripheral processes, such as hyperactivity of primary nociceptors, are critical to the expression of pain after SCI. In adult rats, truncal stimulation within the tuning range of C-low threshold mechanoreceptors (C-LTMRs) contributes to pain hypersensitivity and elevates respiratory rates (RRs) after SCI. This suggests that C-LTMRs, which normally encode pleasant, affiliative touch, undergo plasticity to transmit pain sensation following injury. Because tyrosine hydroxylase (TH) expression is a specific marker of C-LTMRs, in the periphery, here we used Th-Cre adult mice to investigate more specifically the involvement of C-LTMRs in at-level pain after thoracic contusion SCI. Using a modified light-dark chamber conditioned place aversion (CPA) paradigm, we assessed chamber preferences and transitions between chambers at baseline, and in response to mechanical and optogenetic stimulation of C-LTMRs. In parallel, at baseline and select post-surgical timepoints, mice underwent non-contact RR recordings and von Frey assessment of mechanical hypersensitivity. The results showed that SCI mice avoided the chamber associated with C-LTMR stimulation, an effect that was more pronounced with optical stimulation. They also displayed elevated RRs at rest and during CPA training sessions. Importantly, these changes were restricted to chronic post-surgery timepoints, when hindpaw mechanical hypersensitivity was also evident. Together, these results suggest that C-LTMR afferent plasticity, coexisting with a potentially facilitatory state of sympathetic activation, drives at-level affective pain following SCI in adult mice.

**Contribution to the field statement:** Preclinical studies have only recently sought to understand biological mechanisms connecting peripheral afferent neuron hypersensitivity to pain. Using mechanical brush and selective optogenetic activation in transgenic mice, this study investigated the role of an implicated subpopulation of sensory afferents, the C-low threshold mechanoreceptors (C-LTMRs), that may be transformed from pleasurable touch- to pain-encoding neurons after spinal cord injury (SCI). C-LTMR activation after SCI evoked at-level conditioned pain responses associated with sympathetic activation (increased respiratory rates). These studies provide a foundation for research into the peripheral afferent mechanisms of pain that is expected to translate into new options for pain control in humans.

## 1. Introduction

Neuropathic pain, a common type of pain arising from primary damage to the nervous system, is a clinically relevant outcome of spinal cord injury (SCI) that may be experienced at the level of injury (i.e. “at-level” pain) (Finnerup et al., 2001; Siddall et al., 2003; Felix et al., 2007). Despite the prevalence of pain in approximately 70% of SCI patients (Siddall and Loeser, 2001), pharmacological treatments are often inadequate, only slightly reducing pain intensity (Baastrup and Finnerup, 2008). Although prior studies have focused primarily on plasticity within the injured spinal cord, there is increasing recognition that peripheral plasticity contributes to pain after SCI. For example, increased spontaneous activity in primary nociceptors generates pain (Bedi et al., 2010), and peripheral nociceptive input exacerbates pain behavior (Garraway et al., 2014; Martin et al., 2019), after SCI. While these previous studies identify activity in nociceptors as crucial to pain hypersensitivity, the role of specific afferent subpopulations, including non-nociceptors, in SCI-induced neuropathic pain remains poorly understood.

C-low threshold mechanoreceptors (C-LTMRs) may represent one group of primary afferents that contribute to pain after SCI, as they have been implicated in other injury models (Seal et al., 2009; Mahns and Nagi, 2013). C-LTMRs are small diameter, unmyelinated afferents that innervate the trunk hairy skin. They terminate in lamina II of the dorsal horn (Li et al., 2011), projecting onward to wide dynamic range (WDR) lamina I spinoparabrachial projection neurons (Andrew, 2010; Craig, 2010). C-LTMRs are defined by tyrosine hydroxylase (TH) expression (Li et al., 2011; Lou et al., 2013), and normally encode the affective component of pleasurable, affiliative touch (Iggo, 1960; Bessou et al., 1971; Olausson et al., 2002; Loken et al., 2009; Li et al., 2011; Liljencrantz and Olausson, 2014; Zimmerman et al., 2014). In SCI patients with ongoing pain, gentle brush stimuli applied to hairy skin at the lesion level, a stimulus associated with activation of C-tactile fibres (CTs, the equivalent of C-LTMRs in humans) (Loken et al., 2009) produced hyperesthesia and allodynia (Finnerup et al., 2003). Thus, C-LTMRs are an important target for chronic pain research, but few preclinical studies have related SCI-induced pain to cutaneous trunk signaling, including afferent plasticity at the segmental level of injury [e.g. (Oatway et al., 2004; Yezierski et al., 2004; Crown et al., 2006; Bedi et al., 2010)] where these afferents reside. C-LTMRs may activate sympathetic pathways as part of the allodynic response. Conversely, sympathetic nerve stimulation directly excites C-LTMRs and increases their sensitivity to applied mechanical stimuli (Roberts and Levitt, 1982; Roberts and Elardo, 1985; Barasi and Lynn, 1986). Injury-induced pain is associated with increases in sympathetic activity (Kinnman and Levine, 1995b; a; Jänig et al., 1996; Ramer et al., 1999; Zhang et al., 2004), and nociceptive stimuli can activate the sympathetic nervous system, increasing heart rate and respiratory rate (RR) (Wolf and Hardy, 1941; Culman et al., 1997; Loggia et al., 2011; Santuzzi et al., 2013). Slowing RR (Grant and Rainville, 2009; Noble et al., 2017) or blocking sympathetic pathways (Pinheiro Fde et al., 2015) has been shown to mitigate pain sensitivity.

Our laboratory recently found that mechanical stimulation within the tuning range of C-LTMRs, delivered to adult rats after SCI, increased RRs at timepoints consistent with the expression of pain (Noble et al., 2019). SCI rats also had higher basal RRs acutely after injury. These results support the involvement of C-LTMRs in at-level SCI pain and suggest that acute changes in RRs may serve as a physiological index of pain. It is conceivable that C-LTMRs undergo central or peripheral plasticity after SCI. To identify the specific role C-LTMRs play in SCI-induced neuropathic pain, we undertake studies in adult transgenic mice using a novel conditioned place aversion (CPA) paradigm. We use optogenetic techniques to enable selective stimulation of TH-expressing cutaneous afferents in the trunk skin, and study its effects on affective-motivational (deemed ‘affective’) pain behavior and RRs after SCI.

## 2. Materials and Methods

### 2.1 Animal care and generation of transgenic mice

All experiments were performed in adult male and female transgenic mice expressing Cre recombinase under the regulatory control of the mouse tyrosine hydroxylase (*TH*) gene, i.e. ‘TH-Cre’ mice. Mice were approximately 10 weeks old and weighed 20-22 grams (females) and 24-26 grams (males) at commencement of experimental procedures. They were housed in standard cages in a vivarium on a 12:12-hour light-dark cycle (7 am lights on / 7 pm lights off), with all behavioral testing and respiratory recordings performed during the light period. Animals were fed standard rodent diets *ad libitum*. Experimental procedures were approved by the Animal Care and Use Committee of Emory University and conformed to national standards for the care and use of experimental animals and the American Physiological Society’s “Guiding Principles in the Care and Use of Animals”.

Mice bred in our animal colony originated from one of two commercially available strains; Mutant Mouse Resource & Research Centers (MMRRC) #017262 or Jackson Laboratory (JAX) #025614 mice; the latter strain expresses a tamoxifen-inducible Cre recombinase (Cre/ERT2). For optogenetic experiments, the TH-Cre and tamoxifen-inducible ‘TH-Cre-ER’ mice were crossed with a strain expressing a Cre-dependent channelrhodopsin (ChR)-2 – eYFP fusion protein following exposure to Cre recombinase [Ai32(RCL-ChR2(H134R)/eYFP; JAX #024109], generating TH::ChR2-eYFP transgenic mice. This allows for direct optical targeting of C-LTMRs based on their known molecular profile (Li et al., 2011). The tamoxifen-inducible Cre mice crosses of strain TH::ChR2-eYFP were treated with tamoxifen dissolved in peanut oil [(TAM), 5 mg/day for 2 days (1 day apart) or 2 mg/day over 3 consecutive days), by subcutaneous (SC) administration at the scruff of the neck] to induce transgenic expression. TAM was administered at or shortly after weaning, at around 3 weeks of age, under light isoflurane anesthesia. This dosing regimen was chosen based on preliminary electrophysiology and immunohistochemistry experiments (e.g., **Figure 1A**) that provided confirmation of Cre-driven YFP expression.

**Figure 1.**
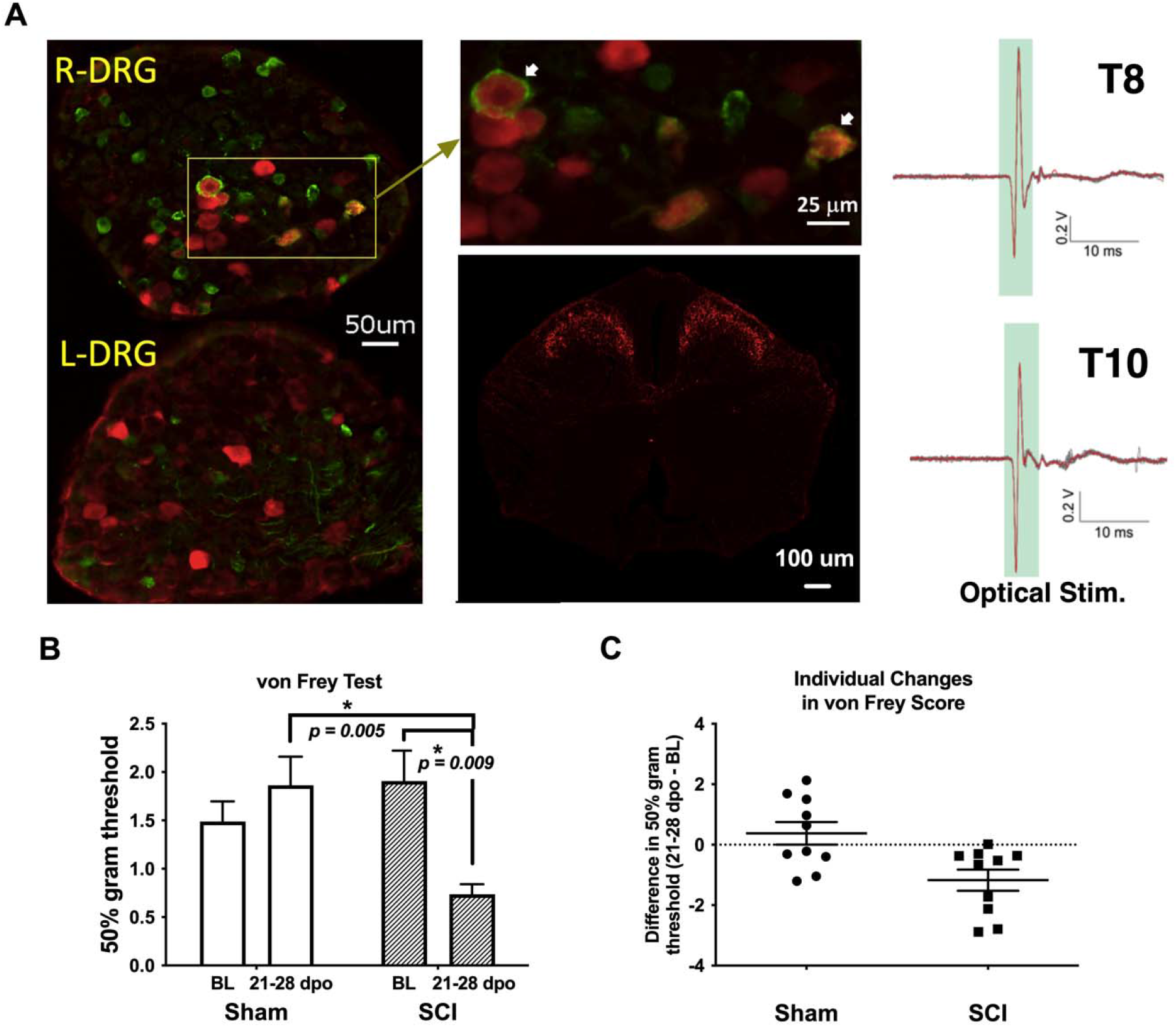
Validation of TH-transgenic mice and SCI model of chronic neuropathic pain. (**A**) *Left and top middle*: Co-expression of YFP and pERK in DRG ipsilateral (R-DRG) but not contralateral (L-DRG) to the side of optical stimulation (note pERK in red and YFP in green). *Bottom middle:* Representative immunohistochemical image of a thoracic (T) level 10 spinal cord section in one naïve TH::tdTomato adult mouse, demonstrating red fluorescent protein indicative of TH expression in superficial laminae of the spinal cord dorsal horn. Observed projection patterns support the recruitment of TH-positive C-LTMRs in the spinal cord (Li et al., 2011). *Right:* Electrophysiological recordings validated successful afferent activation in a TH::ChR-eYFP mouse skin-nerve preparation. Activation was obtained from all nerves following trains of optogenetic stimulation (2.5 V, 5 Hz stim., ten 5-ms pulses). (**B**) Graph shows changes in mechanical reactivity to von Frey stimulation in SCI and sham control mice (N=10 sham, N=10 SCI). SCI mice showed mechanical hypersensitivity at 3-4 weeks post-injury, with significantly reduced hindpaw withdrawal thresholds compared their own baseline values and sham mouse thresholds at the same chronic timepoints. On average, SCI mice withdrawal thresholds were reduced by 61% compared to sham controls. (**C**) Differences in withdrawal threshold between chronic and baseline timepoints are plotted for individual mice with mean values superimposed (Sham and SCI), illustrating the basis of group differences.

### 2.2 Surgical procedures and recovery

Mice were deeply anesthetized with isoflurane (5%, gas; lowered to 2-3% once stable anesthesia was achieved). Under sterile conditions, a skin incision and dorsal laminectomy exposed the underlying spinal cord at lower thoracic level 10 (T10). For midline contusion injuries, mice received a 70 kdyne, zero dwell time, impact onto the dorsal surface of the spinal cord with an Infinite Horizon impactor (IH-0400 Impactor, Precision Systems and Instrumentation, Fairfax Station, VA, USA) as we previously described (Parvin et al., 2021; Martin et al., 2022). Care was taken to ensure that dorsal roots were not damaged by the laminectomy or impact, and on-target bilateral bruising of the dorsal spinal cord was verified by examination under a dissecting microscope. The overlying muscle and skin were sutured and the wound area treated with triple antibiotic ointment (bacitracin-neomycin-polymyxin B) topically. Sham control mice underwent the same surgical procedure but without receiving SCI. All mice (SCI and sham) were left to recover on a heated pad. Animals were given meloxicam (5 mg/kg, SC) immediately after surgery. They were also administered Lactated Ringer’s solution (0.5 mL, intraperitoneally [IP]) immediately after surgery, and an identical injection of 0.9% sterile saline daily for the first 48 hours after surgery, to maintain hydration. Subsequent administration of saline was given only as needed. Mice received the antibiotic Baytril (2.5 mg/kg, SC) immediately after surgery and daily each morning up to 7 days post operation (dpo) to minimize the risk of urinary tract or bladder infection in SCI animals. Experimenters manually expressed mouse bladders twice daily for the duration of experiments or until animals had empty bladders for three consecutive days. Mice were weighed daily for 1 week after surgery and were subsequently weighed weekly and on days when behavioral tests were undertaken. Throughout the experiments, they were carefully monitored for signs of infection or distress. Mice were assessed for impairment of locomotor function at 1 dpo using the Basso Mouse Scale (BMS) (Basso et al., 2006), to ensure effectiveness of the injury. SCI mice were only included in the study if they recorded BMS scores of 0 or 1 at 1 dpo. Three mice (2 SCI and 1 sham) either died before the end of behavioral data collection or were SCI mice presenting with BMS scores greater than 1 at 1 dpo and were therefore excluded from subsequent analysis (also see Results).

### 2.3 Behavioral assays

#### 2.3.1 von Frey Test

At baseline (before surgery) and either 21 or 28 dpo (see below), a subset of mice was transferred to acrylic chambers in a behavioral testing room for the von Frey test of mechanical sensitivity. This assay was not performed 1 day after surgery since SCI mice had BMS scores of 0 or 1, indicating very little hindpaw placement. All mice were acclimated to the behavioral suite and testing apparatuses for at least 3 days prior to surgery. On testing days, following a 30-minute acclimation period, individual animals were assayed for mechanical sensitivity (paw withdrawal responses) according to the established up-down method (Chaplan et al., 1994). Calibrated von Frey hairs (NC12775-99, North Coast Medical, Inc., Morgan Hill, CA, USA) starting with filament evaluator size 3.22 (target force 0.16 grams) were administered from below a metal mesh platform to test each animal’s sensitivity to mechanical stimulation of the hindpaw. Right and left paw withdrawal thresholds were averaged to determine overall mechanical sensitivity. A reduction in von Frey withdrawal threshold values from their baseline levels corresponded to increased mechanical sensitivity.

#### 2.3.2 Conditioned Place Aversion (CPA)

To provide a validated assessment of at-level, C-LTMR-mediated pain after SCI, our laboratory developed a modified CPA paradigm to model affective pain in mice using protocols adapted from previous studies (Hummel et al., 2008; Yang et al., 2014; Bagdas et al., 2016; Refsgaard et al., 2016; Wu et al., 2017) including our own (Martin et al., 2022). The custom-built CPA apparatus consisted of black and white chambers separated by a small partition. Each CPA box contained a small window permitting entry of a brush or fiber optic cable for optogenetic stimulation. Two separate experiments were conducted. The first investigated more ethologically relevant brush stimulation, with results identifying a time window of peak behavioral sensitivity for targeting C-LTMRs with optogenetic techniques in the second experiment. In both experiments, stimuli were delivered across the animal’s trunk for a distance of ~3 cm, and stimulations were provided with an interstimulus interval of 30-60 seconds. Video recordings were collected throughout testing periods, and the percentage of time spent in the non-stimulated chamber before and after stimulation taken to indicate relative place aversion. Note that for brush CPA, all mice were stimulated in the dark chamber, whereas to improve experimental design the initially preferred chamber (occasionally the light chamber) was always used as the stimulation chamber for optical CPA; hence the use of % in light (brush CPA) versus % in non-preferred (optical CPA) chambers as reported outcome variables. For brush CPA, stimulation was undertaken at weekly timepoints to establish a preliminary timeline of C-LTMR-related pain aversion. Subsequently, results from these studies were used to guide selection of timepoints for a 5-day CPA paradigm tailored to peak stimulus aversiveness for optical stimulation in the second experiment (separate mice).

##### 2.3.2.1 Brush Stimulation CPA Cohort

This experiment was done in 14 TH-Cre mice (N=7 each, SCI and sham). Using a modified light-dark chamber CPA paradigm, we assessed preferences at timepoints ranging from 1 day to 5 weeks after surgery. Starting 1 week after surgery, mice were given truncal stimulation with a small histology brush (Camel hair #4, Ted Pella, Inc., Redding, CA, USA) to study pain affect. Each 30-minute test consisted of three distinct periods (compare this to the prolonged nature of optical CPA below, where in order to permit time for greater conditioned responses to develop, once peak affect was defined these three periods were spaced out over 5 days with no more than one occurring on a given day). During the pre-stimulation test, mice were given free access to both chambers (dark and light) for 10 minutes. Then, mice received 5 minutes of mechanical stimulation to the trunk with the brush (once/min), while confined to the dark chamber. For this, we chose a manual stimulation speed (~1 cm/s) that corresponded to the tuning properties of C-LTMRs (Loken et al., 2009; Noble et al., 2019). During a counterbalanced 5-minute ‘control’ epoch in the light chamber, mice received no mechanical stimulation. Finally, during the 10-minute post-stimulation test, mice were again permitted access to both chambers for 10 minutes with no stimulation. Mice in this experiment underwent von Frey testing at baseline and 21 dpo.

##### 2.3.2.2 Optogenetic Stimulation CPA Cohort

This experiment was done in 16 mice as follows: 12 TH-Cre-ER mice (N=7 SCI, N=5 sham) and 4 TH-Cre mice (N=2 SCI, N=2 sham), both ChR crosses. Results were combined for presentation as there were no statistical differences between genotypes, so they are collectively referred to as TH::ChR2-eYFP mice. To encapsulate the period of peak sensitivity observed with brush, centered at 28 dpo (see Results), the optical CPA test ran from 26-30 dpo. For the pre-conditioning test (26 dpo), mice were placed in the light chamber and given free access to the full apparatus for a 30-minute test period to determine baseline time spent in each chamber. They then received daily 30-minute conditioning sessions over the course of three days (27-29 dpo). In their preferred chambers, animals received optical stimulation (once/minute for 10 minutes, after a 5-minute habituation period) provided with a laser to activate C-LTMRs. A fiber optic cable was positioned close to the abdomen of mice and blue light illumination swept across the skin while animals were pseudo-restrained. For pseudo-restraint, mice were moved to an acrylic cylindrical tube (Rodent Restrainer; IITC Life Science, Woodland Hills, CA) that provided them with adequate room to move forwards and backwards but not sideways. This setup balanced the confinement necessary for high-fidelity electric field sensor recordings with stress reduction measures. Mice were acclimated to the cylinder for at least 3 days prior to surgery, and RR was confirmed to stabilize. Fur was shaved at the segmental level of injury for better illumination of cutaneous and subcutaneous nerve fibers. In the control, non-preferred chambers, mice received fake “stimulation” delivered with the laser box powered off but under similar pseudo-restraint (again at once/minute for 10 minutes after a 5-minute habituation). The frequency, duration, and intensity of the light stimulation were manually controlled. The fiber optic cable delivered up to 5,204 mW/cm^2^ of blue light to the skin along the trunk. One day after completion of the conditioning phase (at 30 dpo), each mouse again was allowed free access to explore both chambers for a 30-minute post-conditioning test with no stimuli present. Mice in this experiment underwent hindpaw von Frey testing at baseline and 28 dpo, to verify that hindpaw sensitivity was evident at the time of CPA behavioral assessment.

### 2.4 Respiratory monitoring and analysis

Resting and optically-evoked respiratory measurements were obtained in a subset of mice used in the behavioral assays above. At baseline, and 1 and 21 dpo (with a subset at 7 dpo), mice were sequentially moved to a cylindrical tube (Rodent Restrainer, IITC Life Science, Woodland Hills, CA, USA) for 1-hour respiratory recording sessions to establish basal, spontaneous (resting) RR. In mice receiving optical CPA, the longer duration of training during conditioning sessions (days 2-4 of the 5-day paradigm) as compared to brush CPA also permitted continuous monitoring of evoked RR. In both cases, recordings were obtained using remote sensors developed for this purpose.

#### 2.4.1 Resting RR

Data from the 1-hour recording sessions was used to determine resting RR values for baseline and several post-surgery testing points. Prior to experimentation, non-contact electric field sensors (EPIC, Plessey Semiconductors, Plymouth, UK) were affixed externally to the sides of rodent enclosures, with their wires connected to a power supply box. This box also adapted connections to BNC outputs for subsequent signal digitization and data collection. The analog sensor signal was low-pass filtered (bandwidth DC to 12 Hz) and digitized at unity gain and a sample rate of 1 kHz. The digitized data was continuously output to a Windows computer running LabVIEW (National Instruments, Austin, TX, USA), where recorded data were processed by a customized interface (program designed by William N. Goolsby). All manual and automated analysis of raw sensor output was accomplished using Clampfit analysis software (Molecular Devices, San Jose, CA, USA). In Clampfit, raw signals were analyzed and filtered, and threshold-based detection of individual breaths was performed. Clampfit calculated instantaneous RRs over the entire recording period, which were then averaged over periods of rest to determine a final value for resting RR. The first 20 minutes of each recording period, during animal acclimation, were left unscored. Video recordings verified that final values exclusively captured RR during animal resting states. Although it was attempted, recording RR during von Frey stimulation proved unreliable due to excessive motion artifacts.

#### 2.4.2 Optically-evoked RR

Respiratory recordings were collected non-invasively during CPA procedures (optogenetic and control stimulation) using the remote sensors described above. Similar to resting RR, optically-evoked RR was calculated in Clampfit from scatterplots of instantaneous RR – in this case, concomitant with conditioning sessions in the CPA chamber (27-29 dpo). Approximately 10-second epochs of high-fidelity respiratory recordings were isolated from each interstimulus interval (i.e., between consecutive stimulations). Several replicate measurements of RR were then averaged to obtain individual data points for all six scorable time periods (the three conditioning sessions, with each session comprised of laser stimulation and control conditions).

### 2.5 Fluorescent histochemical and electrophysiological validation

To *i)* confirm that optical stimulation activated TH-positive cells in the dorsal root ganglia (DRG), and *ii)* to validate the expression pattern of TH projections in the spinal cord, fluorescent histochemistry was performed. To *i) validate optical activation of the C-LTMRs*, TH-Cre-ER mice were crossed with channelrhodopsin mice (JAX #024109) and the progeny (TH::ChR2-eYFP mice) were treated with TAM as described above. Two TAM-treated TH::ChR2-eYFP SCI mice were optically stimulated for 10 minutes in an identical manner to CPA conditioning sessions above to determine the effect of stimulation treatment on markers of neuronal activation. Mice were perfused 30 minutes following stimulation for immunolabeling, with cellular co-labeling of phosphorylated ERK (pERK) protein and TH (C-LTMRs, represented by YFP) shown in the DRG (**Figure 1A,** left and top middle). pERK is a key marker of neuronal activity and pain hypersensitivity (Xu et al., 2008; Gao and Ji, 2009; Garraway et al., 2011). To *ii) reveal the expression pattern of TH projections in the spinal cord*, TH-Cre-ER mice were crossed with tdTomato mice (JAX #007908) and the progeny (TH::tdTomato mice) were treated with TAM as described above. Two naïve TH::tdTomato mice were imaged (**Figure 1A,** bottom middle).

In both cases, mice were anesthetized with urethane (1.2 g/kg, IP) and transcardially perfused with phosphate-buffered saline (PBS) followed by 4% paraformaldehyde (PFA) in PBS. The spinal cord and/or DRGs were dissected and post-fixed in 4% PFA for 2 hours before being transferred to 30% sucrose for cryoprotection. Transverse sections (20 μm) of lower thoracic cord or DRG were cut with a cryostat (Leica Microsystems, Buffalo Grove, IL, USA) and mounted on Superfrost Plus slides (Fisher Scientific, Pittsburgh, PA, USA). The slides were dried overnight and then stored at −80°C until the time of use. For fluorescent immunohistochemistry, the spinal cord and DRG sections were washed in PBS and PBS with 0.1% Triton X-100 (PBS-T), then incubated in blocking solution (5% donkey serum [#017-000-121, Jackson ImmunoResearch Laboratories, Inc., West Grove, PA, USA] in PBS-T) for 1 hour at room temperature. To *i) validate optical activation of the C-LTMRs*, sections were then incubated in rabbit anti-pERK (1:200; #4370-Cell Signaling, Inc., Lake Placid, NY, USA) and/or chicken anti-green fluorescent protein (GFP) (1:500; #ab13970—Abcam, Inc., Cambridge, UK) primary antibody in blocking solution for 24 hours at room temperature in a humid chamber on a gentle rotator plate (GFP antibody also detects YFP). The sections were washed in PBS-T and then incubated in Cy3-conjugated donkey anti-rabbit (1:250; #711-165-152-Jackson ImmunoResearch Laboratories, Inc., West Grove, PA, USA) and/or Cy2-conjugated donkey anti-chicken (1:100; #703-225-155—Jackson ImmunoResearch Laboratories, Inc., West Grove, PA, USA) fluorescent secondary antibody in blocking solution for 1 hour at room temperature. To *ii) reveal the expression pattern of TH projections in the spinal cord*, sections were instead incubated in rabbit anti-red fluorescent protein (RFP) (1:250 in 1% BSA-PBS; #600-401-379— Rockland Immunochemicals, Inc., Gilbertsville, PA, USA) primary antibody in blocking solution for 48 hours at 4°C in a humid chamber on a gentle rotator plate. The sections were washed in PBS-T and then incubated in Cy3-conjugated donkey anti-rabbit (1:500 in 1% BSA-PBS; #711-165-152—Jackson ImmunoResearch Laboratories, Inc., West Grove, PA, USA) fluorescent secondary antibody in blocking solution for 1 hour at room temperature.

Following another series of washes in PBS, all sections were mounted in ProLong Gold anti-fading mounting medium (Invitrogen, Eugene, OR, USA), and coverslipped. Serial images were taken with a digital/confocal microscope (Keyence VHX-7000 Digital Light Microscope, 4X or 20X objective lens, Osaka, Japan) and stitched together to produce conglomerate images using Olympus Fluoview version 5 (Olympus America Inc., Center Valley, PA, USA).

Details of electrophysiological procedures have been published (Provost, 2019). Briefly, electrophysiological recordings were undertaken to validate optogenetic C-LTMR activation in adult SCI mice ~3 months post-injury using a TH::ChR-eYFP mouse skin-nerve preparation. Recordings were obtained from dorsal cutaneous nerves at thoracic spinal segments T8 through T12. The skin, with attached dorsal cutaneous nerves, was placed epidermal-side down over a hole in a recording dish and held in place by a cap and screws. The skin formed a seal and the dish was filled with a circulating bath maintained at 26°C. The distal end of the nerve was attached to a tight-fitting suction electrode, and the dish was placed over a circular hole on an elevated platform. The fiber optic cable was affixed to a computer-controlled robotic arm on the underside of the platform. Fiberoptic blue light pulses were used to evoke activity in TH-positive C-LTMRs. Activation was obtained from nerves following trains of optogenetic stimulation (2.5 V, 5 Hz stim., ten 5-ms pulses).

### 2.6 Statistical analysis and blinding

CPA and von Frey behavioral assessments were performed by the same individual. Although it was impossible to blind experimenters to animal group due to obvious hindlimb impairment in SCI mice, adequate steps were undertaken to minimize experimental bias. For instance, RR recordings and CPA videos were coded prior to scoring. This ensured blinding of RR data since recordings were scored independent of animal observation. Also, although CPA scoring necessitated observation of animal behavior, the simple nature of these measures (time in each chamber and crossings between chambers) left little room for experimenter bias.

Data are presented as mean ± SEM, unless otherwise noted. All behavioral (von Frey and CPA tests) and RR (resting and optically-evoked RR) data were analyzed independently for each group (sham or contusion SCI) using one-way repeated measures (RM) ANOVA with days post-operation as the within-subjects factor. Post-hoc tests corrected for multiple comparisons were performed in the case of significant results. In all cases, the choice of multiple comparisons test was selected according to GraphPad Prism (taking appropriateness for the particular dataset into account). Paired t-tests were occasionally performed as planned comparisons, as indicated in the text. Comparison between groups was accomplished using two-way RM ANOVA with days post-operation as the within-subjects factor and group (sham or SCI) as the between-subjects factor. As above, post-hoc tests corrected for multiple comparisons were performed in the case of significant results. Unpaired t-tests comparing group means were occasionally performed as planned comparisons, again as indicated in the text. To quantify the relationship between affective (CPA) pain, hindpaw mechanical sensitivity, and changes in RR, correlation analyses were undertaken at several different timepoints corresponding to observed changes in SCI mice. Statistics were performed with GraphPad Prism (GraphPad Software, La Jolla, CA, USA), with significance set at p < .05 and two-tailed tests.

## 3. Results

Prior to the two primary experiments below, mice were assigned to receive sham surgery (N=15; 8 females and 7 males) or thoracic level (T10) moderate midline contusion SCI (N=18; 10 females and 8 males). One sham mouse and two SCI mice met exclusion criteria (see Materials and Methods) and were not included in subsequent analysis. Two TH::ChR2-eYFP SCI and two naïve TH::tdTomato mice were also used for immunofluorescence. Data from male and female mice were combined as there were no statistical differences between sexes for any of the primary outcome measures assessed.

### 3.1 Fluorescent Histology

A representative image of pERK and YFP double-labeling in DRGs following optical stimulation is shown in **Figure 1A** (left and top middle). Consistent with a previous report (Li et al., 2011), our preliminary fluorescent histology revealed RFP labelling indicative of TH expression in the superficial laminae of the thoracic spinal cord dorsal horn as shown in **Figure 1A** (bottom middle). The similarity of observed projection patterns to those previously reported suggests that we are identifying afferents whose cellular and anatomical properties are consistent with those of C-LTMRs. Selective recruitment of C-LTMRs in our study is further supported by *ex-vivo* electrophysiological recordings validating activation of C-LTMRs by optical stimulation in some of the same TH::ChR-eYFP mice used for behavioral experiments, shown in **Figure 1A** (right).

### 3.2 von Frey Test

Sham mice did not develop any differences in sensitivity in the von Frey test from baseline to 21-28 dpo (*t_(9)_* = 1.0, *p* > .05, paired t-test). In contrast, SCI mice scored for hindpaw sensitivity at these same chronic post-injury timepoints clearly developed mechanical hypersensitivity (*t_(9)_* = 3.4, *p* < .01, paired t-test), representing a 61.4% reduction in 50% gram threshold from baseline values (**Figure 1B, C**). SCI mice were also significantly more sensitive than their sham counterparts at the chronic timepoint (*t_(18)_* = 3.6, *p* < .005, unpaired t-test). The separate cohorts of SCI mice scored at 21 and 28 dpo presented indistinguishable mechanical thresholds by the end of testing (*t_(8)_* = 0.03, *p* > .05, unpaired t-test) with mean 50% gram thresholds of 0.74 and 0.73, respectively, so data were pooled to increase statistical power.

### 3.3 Conditioned Place Aversion (CPA)

A light-dark box CPA paradigm was used to assess affective pain responses (contextual aversion to C-LTMR stimulation by brush or laser). Chamber transitions and time spent in each chamber were monitored as depicted in **Figure 2A**.

**Figure 2.**
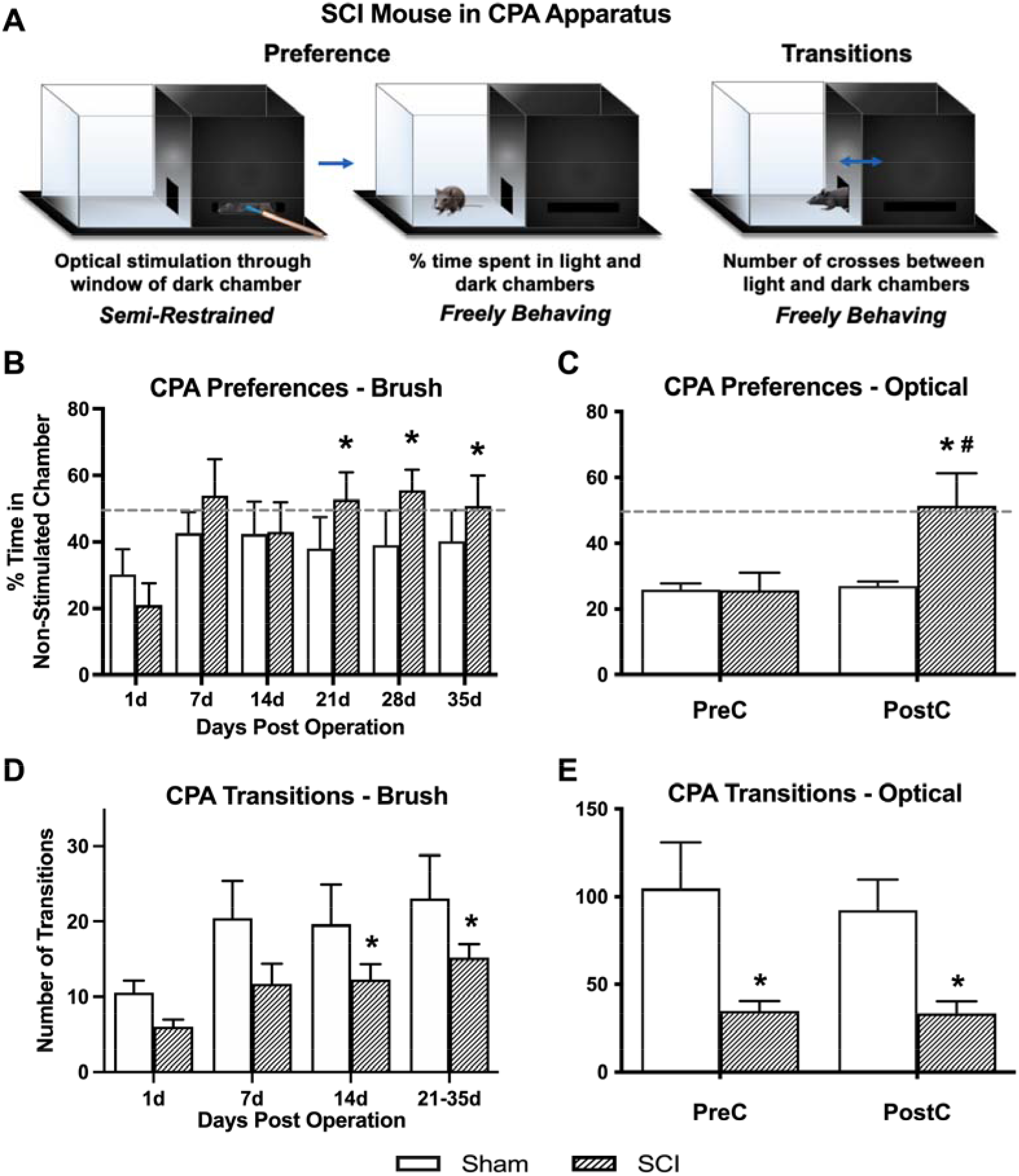
CPA affective pain responses following mechanical and optical C-LTMR stimulation. **(A)** Schematic depicting the two-chamber (light-dark) CPA apparatus, illustrating mouse positioning during stimulation (*left, optical stimulation shown*) and free exploration in the light chamber (*middle*) or transitioning between chambers (*right*). **(B)** SCI mice administered mechanical brush stimulation while confined to the dark chamber later developed a preference for the light “escape” chamber that peaked 4 weeks after injury (*\*p<0.05 versus 1 dpo*). **(C)** We found similar changes in preference following selective optical stimulation of TH-expressing sensory afferents, the C-LTMRs (*\*p<0.05 SCI pre-conditioning [PreC] versus post-conditioning [PostC]; #p<0.05 SCI versus Sham at PostC*). For (B) and (C), horizontal dotted lines indicate 50% time spent in non-stimulated chamber. (**D,E**) The total number of side-to-side transitions during CPA behavioral experiments was used as a broad assessment of locomotor activity, revealing the expected impairment after injury in SCI mice. Note that the longer duration of CPA pre- and post-stimulation sessions in the second experiment (optical stimulation) resulted in a greater overall number of side-to-side transitions compared to brush stimulation. For (B) and (D), N=7 mice per group; for (C) and (E), N=7 sham and N=9 SCI.

#### 3.3.1 Brush CPA – % time in light chamber

Overall averages combining the pre- and post-stimulation periods were used to derive our principal conclusions in regard to the aversiveness of C-LTMR-targeting brush stimulation since these values represent a more robust average (over 20-minute weekly periods) of an animal’s total preference for escaping the brush-associated context. Values for % time in the light chamber of the CPA apparatus (hereafter referred to as ‘% in light’) at 1 dpo averaged 25.6 ± 5.0% and were statistically indistinguishable between groups (sham: 30.3 ± 7.5, SCI: 21.0 ± 6.6). One-way RM ANOVA within SCI mice revealed a significant increase in % in light over days post-surgery (*F*_(2.0, 11.8)_ = 5.1, *p* < .05). Comparisons between post-surgical ‘baselines’ (at 1 dpo) and subsequent timepoints using Dunnett’s multiple comparisons tests revealed a significant increase in preference for the light chamber at all three chronic timepoints (21 dpo: 52.8 ± 8.2%, *p* < .01; 28 dpo: 55.5 ± 6.3%, *p* < .005; and 35 dpo: 50.9 ± 9.1%, *p* < .05), but not at acute or subacute timepoints (7 or 14 dpo) (**Figure 2B**). In contrast, sham mice did not show changes in preference at any timepoint (*F*_(1.6, 9.6)_ = 0.42, *p* > .05, one-way RM ANOVA), and they never spent more than 42.6% of their time in the light chamber (at 7 dpo), with values even decreasing slightly at chronic timepoints (average from 21-35 dpo: 39.1%). Comparing the two groups using two-way RM ANOVA, there was a significant effect of days post-surgery on % in light (*F*_(2.6, 31.0)_ = 3.5, *p* < .05). However, there was no significant group effect (*F*_(1, 12)_ = 0.66, *p* > .05).

#### 3.3.2 Optogenetic CPA – % time in non-stimulated chamber

To minimize potential ceiling effects, optogenetic experiments provided stimulation to mice in their preferred chamber (occasionally the light chamber) during conditioning sessions. Pre-conditioning values (pre-stimulation at 26 dpo) were similar to those observed in the brush CPA experiment, with both groups spending less than 30% of their time in the non-preferred chamber (sham: 25.9 ± 1.8%, SCI: 25.7 ± 5.5%). In contrast, SCI mice developed a large and significant preference for the non-stimulated chamber following three days of optical stimulation targeting C-LTMRs (51.4 ± 9.9%; *t(_8_)* = 3.4, *p* < .01, paired t-test), while sham mice showed no change in chamber preference (26.9 ± 1.4%; *t_(6)_* = 0.5, *p* > .05, paired t-test) (**Figure 2C**). On average, SCI mice spent 90.9% more time in the non-stimulated chamber during the post-stimulation period than sham controls. Despite large intraindividual variability – especially in SCI mice – the two groups were significantly different (*t_(14)_* = 2.2, *p* < .05, unpaired t-test).

#### 3.3.3 Transitions (Brush and Optogenetic CPA)

Locomotor assessment with the BMS was only undertaken at 1 dpo to confirm injury severity. However, transitions between chambers (i.e. side-to-side crosses) in the CPA paradigm were used as a broad quantification of locomotor function and development of conditioned aversion to targeted C-LTMR stimulation at later timepoints. <u>Brush Cohort:</u> Using two-way RM ANOVA, there was no main effect of group (SCI versus sham) on overall transitions despite a trend (*F*_(1, 12)_ = 2.8, *p* = .12); however, in the same test there was a main effect of dpo (*F*_(1.7, 20.7)_ = 8.5, *p* < .005). There was a significant change in the number of transitions over days post-injury in SCI mice (*F*_(1.5, 9.2)_ = 9.8, *p* < .01, one-way RM ANOVA), with increased transitions at 14 dpo (*p* < .05) and 21-35 dpo (*p* < .005) as compared to 1 dpo using Dunnett’s multiple comparisons tests. Overall sham mouse transitions showed a trend toward varying with dpo (*F*_(1.5, 8.8)_ = 3.5, *p* = .09, one-way RM ANOVA). The increase in transitions from 1 dpo to chronic timepoints within SCI mice but not sham controls supported a gradual recovery of locomotor function post-injury (but not full recovery; see **Figure 2D, E**). <u>Optogenetics Cohort:</u> SCI mice performed significantly fewer transitions between the light and dark chambers than sham mice, during both the pre-stimulation test (sham: 104.7 ± 26.3, SCI: 34.8 ± 5.8; *t_(14)_* = 2.9, *p* < .05, unpaired t-test) and post-stimulation test (sham: 92.3 ± 17.5, SCI: 33.4 ± 7.0; *t_(14)_* = 3.4, *p* < .005, unpaired t-test). There was no significant change in the number of transitions from pre- to post-stimulation in either the SCI (*t_(8)_* = 0.2, *p* > .05, paired t-test) or sham (*t(6)* = 1.2, *p* > .05, paired t-test) mice.

### 3.4 Resting and Evoked RR

As depicted in **Figure 3A**, resting RR was monitored at baseline and several post-surgical timepoints as a proxy for sympathetic activation. The extended duration of optical CPA conditioning sessions also permitted recording of evoked RRs at a chronic timepoint (27-29 dpo).

**Figure 3.**
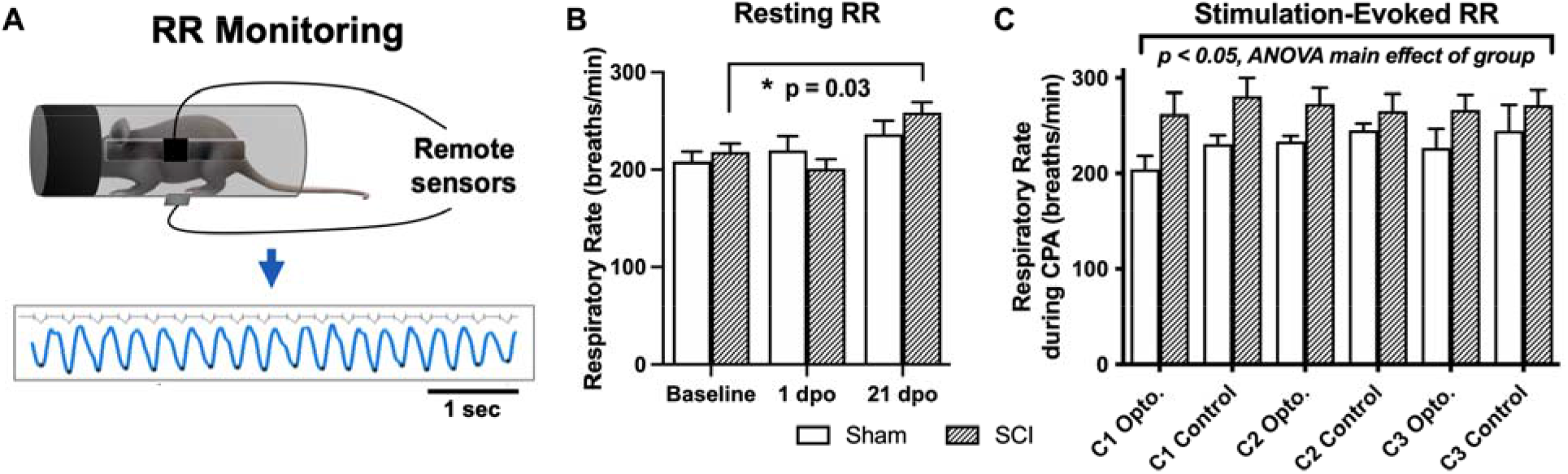
Resting and evoked RR changes in SCI mice. (**A**) Schematic showing respiratory monitoring with remote electric field sensors affixed to an acrylic cylindrical tube. Sensors were positioned facing a small horizontal window in the tube to allow unimpeded RR recordings in semi-restrained mice. Threshold-based detection was performed on individual records to isolate each breath and quantify RRs. (**B**) SCI mice receiving brush or optical stimulation CPA underwent a significant elevation in resting RR at 21 days following injury compared to baseline values, while RRs at acute timepoints (1 dpo and 7 dpo *[subgroup not shown])* remained unchanged. N=7 sham, N=13 SCI (**C**) SCI mice average RRs (271 ±15 breaths/minute) monitored during the optical CPA paradigm were elevated overall compared to sham control average RRs (225 ±16 breaths/minute), and this was significant when data were analyzed over conditioning sessions (*C1, C2, C3*, two-way ANOVA, main effect of group); however, individual stimulations did not appear to increase RR (Šídák’s multiple comparisons tests, sham vs. SCI or opto. vs. control) suggesting that optical stimulation did not acutely increase sympathetic activity. N=4 sham, N=8 SCI

#### 3.4.1 Resting RR

Although there was no main effect of group (SCI versus sham) via two-way RM ANOVA comparing all mice that underwent RR monitoring at baseline, and 1 and 21 dpo (*F*_(1, 18)_ = 0.16, *p* > .05), SCI mice showed a significant elevation in spontaneous (resting) RR over time post-injury (*F*_(1.8, 21.8)_ = 8.6, *p* < .005, one-way RM ANOVA). Planned comparisons revealed a significant elevation in resting RR in SCI mice at 21 dpo compared to baseline (*t_(12)_* = 2.5, *p* < .05, paired t-test) but not compared to sham mice (*t_(18)_* = 1.2, *p* > .05, unpaired t-test). The magnitude of the increase from baseline was substantial, as RR increased from 218 ±9 to 258 ±11 breaths/minute on average (a 40 breaths/minute increase) at this chronic timepoint (**Figure 3B**). In contrast, SCI mice showed no change in resting RR at 1 dpo compared to baseline (*t_(12)_* = 1.3, *p* > .05, paired t-test) or compared to sham mice (*t_(18)_* = 1.1, *p* > .05, unpaired t-test). The subset of SCI mice tested at an acute post-injury timepoint (7 dpo) did not develop a significant increase in resting RR (*t_(4)_* = 1.6, *p* > .05, paired t-test), although the average increase from 216 to 257 breaths/minute suggests this limited dataset may have been statistically underpowered.

#### 3.4.2 Optically-evoked RR

SCI mice undergoing the 5-day CPA paradigm had a higher overall RR than sham controls over conditioning sessions 1-3 (*F*_(1, 53)_ = 9.1, *p* < .005, two-way RM ANOVA). However, when group averages were taken across all conditioning sessions, the difference between SCI and sham mice did not reach significance (*t_(10)_* = 1.9, *p* = .09, unpaired t-test; **Figure 3C**). RR was also not consistently greater immediately following optogenetic versus control stimulation in injured mice (*t_(7)_* = 0.34, *p* > .05, paired t-test) or sham controls (*t_(3)_* = 1.2, *p* > .05, paired t-test), suggesting a dissociation between optically-evoked RR and the changes in preference observed above.

### 3.5 Correlations Between RR Increases and Affective Behavior After SCI

For brush stimulation CPA, there was a significant correlation between pre-stimulation chamber preferences and baseline resting RRs in SCI mice, with higher RRs predicting a preference for the light chamber immediately after injury (r = 0.89, p < .05). Conversely, the same baseline RRs negatively correlated with chamber preference at 7 dpo, reflecting less aversion to the stimulated (dark) chamber following the first occurrence of mechanical truncal stimulation (r = −0.92, p < .05). Baseline resting RRs also negatively predicted changes in chamber preference at 28 dpo (r = −0.88, p = .05). In contrast, there was a negative relationship between changes in resting RR from baseline to 7 dpo and baseline CPA preference (r = −0.87, p = .05), and a positive relationship between 7 dpo RR changes and CPA preferences at the occasion of the first stimulation (7 dpo; r = 0.88, p = .05). The same changes in resting RR at 7 dpo positively predicted changes in chamber preference at 28 dpo (r = 0.96, p < .05; **Figure 4A**). In mice receiving optical stimulation CPA, resting RRs at 1 dpo negatively predicted CPA preferences for the non-stimulated chamber post-stimulation (r = −0.76, p < .05; **Figure 4B**). The same mice with higher acute RRs at 1 dpo also tended to have a clearer chamber preference at baseline, showing a tendency to avoid the aversive (non-preferred) chamber prior to stimulation (r = −0.67, p = .07). Resting RR at 21 dpo was strongly positively correlated with optically-evoked RR responses during CPA conditioning sessions (r = 0.88, p < .005), as were changes in both of these values from baseline RRs (r = 0.94, p < .001; **Figure 4C**). Across both contusion SCI groups, baseline resting RR negatively predicted changes in RR following SCI at the two post-injury timepoints assessed in both experiments, 1 dpo (r = −0.73, p < .005) and 21 dpo (r = −0.76, p < .005). That is, lower starting RRs were associated with greater RR changes experienced acutely and chronically after injury (**Figure 4D**), whereas changes in resting RR from baseline to 1 dpo positively correlated with changes from baseline to 21 dpo (r = 0.66, p < .05). These effects were also observed within the brush and optical CPA groups, when analyzed separately.

**Figure 4.**
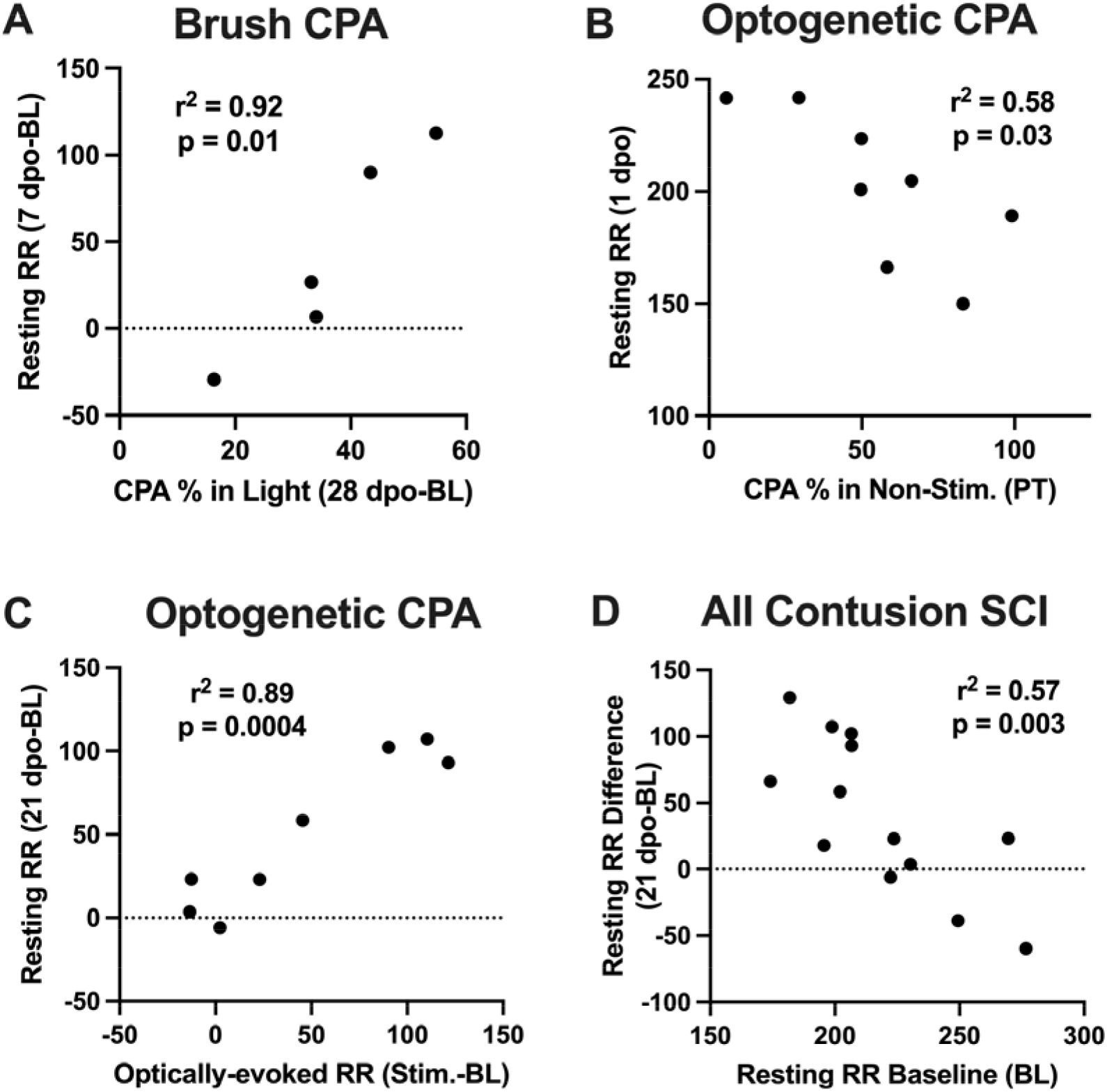
Correlations between RR changes and affective behavior. (**A**) Changes in resting RR at 7 dpo (a sign of acute respiratory plasticity after SCI) positively predicted changes in chamber preference at 28 dpo in the brush CPA cohort. (**B**) In mice receiving optical CPA, resting RRs at 1 dpo significantly predicted CPA chamber preferences post-stimulation, such that mice with higher RRs acutely after injury tended to spend a reduced percentage of their time in the CPA “escape” chamber following stimulation. (**C**) Changes in resting RRs at 21 dpo were strongly correlated with optically-evoked RR responses during CPA conditioning sessions in this same cohort. (**D**) Across SCI mice in both cohorts, there was a negative correlation between baseline resting RRs and injury-induced changes in RR at 1 dpo and 21 dpo, such that mice with the lowest RRs immediately before surgery underwent the greatest RR increases after injury.

## 4. Discussion

This study investigated the contribution of primary afferent plasticity to neuropathic pain following SCI, focusing on C-LTMRs. We found that mechanical stimulation, and optical stimulation of TH-expressing afferents, presumably C-LTMRs, in the trunk skin induced affective pain behaviors in adult mice after SCI. We also found that SCI mice showed a significant elevation of RR at rest and during CPA sessions. All of these responses were seen at chronic timepoints, when hindpaw mechanical hypersensitivity was also evident. These results implicate C-LTMR afferent plasticity in neuropathic pain at or near the level of injury.

It has previously been shown that hyperactivity in nociceptors facilitates pain after SCI (Bedi et al., 2010; Garraway et al., 2014; Martin et al., 2019). However, whether normally non-pain transducing peripheral afferents, the C-LTMRs, can similarly drive neuropathic pain after SCI had not been shown. Expanding on a previous study (Noble et al., 2019), we now provide evidence supporting C-LTMR contributions to evoked, affective pain after SCI. These results also support the proposition that afferent plasticity is critical to the development of affective pain after SCI.

The exact mechanisms by which C-LTMRs might engage nociceptive pathways after SCI have not been elucidated in the current study, although several possibilities can be proposed. The transformation of C-LTMR responses following injury could result from changes in the intensity of C-LTMR activation and/or postsynaptic responsiveness to C-LTMR activity; for example, less C-LTMR activation or downstream responsiveness to C-LTMR activity may elicit pleasant feelings while more activity or heightened dorsal horn responsiveness after SCI may elicit pain. It is also conceivable that C-LTMRs adopt a nociceptor-like phenotype after SCI and thereby show an increase in spontaneous firing (Bedi et al., 2010), a hallmark of SCI-induced pain hypersensitivity. An alternative possibility is that C-LTMRs, like other classes of small diameter primary afferents, undergo anatomical reorganization at their central terminals (e.g. (Weaver et al., 2001; Detloff et al., 2014)). Central sprouting of C-LTMRs might also cause them to project to deeper dorsal horn laminae where they can in turn relay sensory information via nociceptive specific and WDR second-order neurons. Afferent sprouting, including of sympathetic fibers, has also been seen in the periphery and DRG after nerve injury (e.g. (Ramer and Bisby, 1997; Ruocco et al., 2000)), and this is believed to play a role in injury-induced pain (Kinnman and Levine, 1995b; a; Jänig et al., 1996; Ramer et al., 1999; Zhang et al., 2004). Afferent sprouting with concomitant increases in nociceptor receptive fields has been shown to accompany nerve growth factor-induced hyperalgesia (Pertens et al., 1999). Although the exact locus and type of C-LTMR plasticity that occurs after SCI is still unknown, we can hypothesize based on the aforementioned studies that C-LTMRs are capable of undergoing plasticity at their cell bodies (increased excitability), peripheral terminals (changes in receptive field and tuning/recruitment properties) or central terminals (afferent sprouting); and that this plasticity is critical to the expression of at-level pain hypersensitivity after SCI.

C-LTMRs have been implicated in both anti-nociceptive and nociceptive processes. Rodent models revealed a C-LTMR-specific inhibitory pathway for long-lasting analgesia, which is activated by the release of TAFA4, a chemokine-like protein found only on these afferents (Delfini et al., 2013; Kambrun et al., 2018). Slow, pleasant brushing at a velocity (3 cm/s) optimal for activation of unmyelinated CTs can also reduce heat pain (Liljencrantz et al., 2017). C-LTMRs are responsive to massage-like stroking of hairy skin *in vivo* and pharmacogenetic activation in freely behaving mice promotes conditioned place preference (Vrontou et al., 2013). However, C-LTMR activity has also been associated with injury-induced allodynia in both mice (Seal et al., 2009) and humans (Liljencrantz et al., 2013; Mahns and Nagi, 2013; Nagi and Mahns, 2013), and pleasant brushing can evoke allodynia in experimental models of pain including hypertonic saline infusion (Nagi et al., 2011). One potential concern in our study is the lack of an increased preference for the CPA chamber associated with C-LTMR stimulation in sham mice, given the reported pleasurable effects of activating these afferents in both mice and humans (Loken et al., 2009; Vrontou et al., 2013). However, since animals were typically stimulated in their preferred chamber (where they spent ~70-80% of their time), it is likely that a ceiling effect precluded observation of any pleasant or rewarding effects of C-LTMR stimulation. Future studies using our model will stimulate a subpopulation of sham and SCI mice in their non-preferred chambers to assess the true polarity of C-LTMR phenotypes. Clearly, additional studies are also needed to elucidate the exact mechanisms that underlie the sufficiency of C-LTMR stimulation for eliciting pain after SCI. For instance, studies combining optogenetic and behavioral approaches with electrophysiological techniques would be valuable. Incorporation of an *ex-vivo* skin-nerve preparation could provide information on changes in C-LTMR recruitment properties and receptive field maps, while an isolated DRG neuron preparation could be used to investigate the emergence of spontaneous activity in C-LTMRs (Bedi et al., 2010), after SCI.

Another key observation noted in this study is the relationship between changes in resting RR and affective pain behavior in SCI subjects. Resting RR was increased at 21 dpo, a finding that paralleled effects observed in adult rats at earlier timepoints (Teng et al., 1999; Teng et al., 2003; Noble et al., 2019). We also found that RR was acutely increased during the same optical C-LTMR stimulation that successfully evoked a pain response. Respiratory complications frequently accompany SCI in humans (Jackson and Groomes, 1994). Because sensory and autonomic systems interact, high RR may be a proxy for the deleterious effects of sympathetic overactivation on maladaptive pain after SCI. This notion is supported by previous studies showing that nociceptive stimuli activate the sympathetic nervous system, thereby resulting in increased heart rate and RR (Wolf and Hardy, 1941; Culman et al., 1997; Loggia et al., 2011; Santuzzi et al., 2013). Building on these reports, we now show that after lower thoracic SCI, touch-transducing primary afferents promote pain hypersensitivity and may concurrently increase RRs. Surprisingly, correlation analyses revealed that while changes in resting RR predicted aversion to at-level brush stimulation after SCI, this same effect did not hold true for C-LTMR-specific optogenetic stimulation. Therefore, it appears that complex relations between RRs and affective pain after SCI are not mediated exclusively by C-LTMRs, but instead may involve multiple cutaneous afferent subpopulations, and/or central circuits. The most likely possibility is that, while C-LTMRs contribute to affective pain after SCI, the mechanisms linking SCI pain to RR changes are largely C-LTMR-independent or depend on a multitude of factors. This is supported by a previous study, which demonstrated a correlation between RRs and paw withdrawal thresholds (i.e., C-LTMR-independent pain) in the von Frey test (Noble et al., 2019).

After peripheral nerve injury or inflammation, sympathetic stimulation excites nociceptors (Habler et al., 1987; Hu and Zhu, 1989; Sato and Perl, 1991; Sato et al., 1993), and norepinephrine can activate nociceptors after injury (Hu et al., 2000; Tanimoto et al., 2011). Our observed aversive effects of optogenetic stimulation after SCI could plausibly be produced by activation of sympathetic postganglionic neurons that excite nociceptors already sensitized by SCI, with the resulting nociceptor (rather than C-LTMR) activity promoting CPA behavior. While our study design cannot rule out this indirect mechanism, immunohistochemistry in SCI mice provided evidence of direct activation of C-LTMRs via optical stimulation (i.e., TH-positive DRG neurons ipsilateral to the dermatomal field of optical stimulation were selectively recruited; **Figure 1A,** left and top right). This supports the specificity of our stimulation paradigm for C-LTMRs, as does their known molecular composition and spinal innervation pattern (Li et al., 2011) (**Figure 1A,** bottom right). Sympathetic postganglionic axons may sprout in the DRG following peripheral nerve injury to enhance sensory neuron activation or even underlie synchronized cluster firing; however, this functional coupling appears to be diffusion-based rather than via direct connections with soma membrane (Chung et al., 1996; Shinder et al., 1999; Zheng et al., 2022). Future studies should clarify the relationship between primary afferents and sympathetic efferents as well as their relative necessity or sufficiency for initiating autonomic motor and behavioral responses after SCI. Nevertheless, we acknowledge that there is a need for caution in interpreting our results until future mechanistic studies are performed to definitively investigate a sympathetic contribution.

The present study introduces a novel approach to assess at-level SCI pain. Although at- and below-level pain is reported by patients with SCI, finding appropriate research models to assess at-level, non-reflexive pain has been challenging. In fact, very few laboratory methodologies have been implemented (Christensen and Hulsebosch, 1997). In addition to demonstrating that RR is a measurable index of at-level pain (Noble et al., 2019), here, we designed a minimally stressful, non-invasive tool to show that truncal stimulation, at dermatomes surrounding the lesion, induced a CPA response. This strategy was especially important since it enabled us to assess C-LTMRs based on their presence in hairy, but not glabrous, skin (Lou et al., 2013). In addition, we show for the first time that supraspinal, non-reflexive pain responses are evoked by stimulation of C-LTMRs following SCI, at a time when mice also display hindpaw mechanical hypersensitivity. Together, these findings confirm the expression of at- and below-level pain after SCI (Finnerup et al., 2001; Siddall et al., 2003; Felix et al., 2007). Mechanistically, dorsal horn damage after injury could lead to deficits in C-LTMR sensory processing and impair input to supraspinal brain centers, including those involved in the modulation of pain affect. C-LTMR dysregulation following dorsal horn injury may also increase pain aversion via facilitated afferent activation, resulting in reduced C-LTMR-mediated hedonic touch processing and a loss of normal analgesic function (Liljencrantz and Olausson, 2014). Plasticity in pain affect and RR responses after SCI could have different underlying pathways (e.g., see (Teng et al., 1999; Teng et al., 2003)).

A notable limitation of our study is the observed difference in magnitude of conditioned aversion evoked by optical versus mechanical C-LTMR stimulation. Several explanations based on our experimental design might account for the difference. First, the increased frequency of testing in mice receiving brush stimulation (once per week versus over one five-day period) could account for the weaker effects observed with this stimulus. Such an occurrence could constitute either habituation to repeated exposure to a stimulus, or alternatively sensitization to the experimental context. The latter possibility could explain the greater behavioral aversion that developed in sham control mice receiving brush versus those receiving optical stimulation, potentially obscuring group differences for brush. Second, it is likely that the reduced length of pre- and post-stimulation sessions in mechanical brush experiments (10 minutes) versus later optogenetic experiments tailored to the timeline of brush outcomes (30 minutes) was a key contributing factor. Brush stimulation, despite being tuned to activate C-LTMRs, will undoubtedly activate other sensory afferent subpopulations as well and is thus non-selective. Therefore, a third possibility is that broadly-administered brush stimulation might mask the selective effects of C-LTMR stimulation by activating cutaneous afferent subpopulations that oppose or neutralize the impact of C-LTMRs. Using a CPA paradigm similar to ours, Chamessian et al. recently found no impact of cutaneous Aβ-LTMR stimulation on aversive behavior (Chamessian et al., 2019).

The present study demonstrates that cutaneous mechanoreceptors can drive pain behavior after SCI. While there is previous evidence suggesting that non-nociceptor Aß afferents can undergo a phenotypic switch to contribute to inflammatory hypersensitivity (Neumann et al., 1996), a similar change in the properties of mechanosensitive non-nociceptors in chronic SCI pain has not yet been shown. Although we have established the sufficiency of C-LTMR stimulation for inducing an aversive CPA response, future studies should optically inhibit C-LTMRs to establish necessity (e.g. in TH-crossed, archaerhodopsin-expressing mice), and thereby fully assess causality. Studies should also investigate whether pain-associated neural responses are attenuated in these transgenic models.

Our results strongly support recent studies demonstrating transformation of C-LTMRs into pain-transmitting afferents after injury (Seal et al., 2009; Mahns and Nagi, 2013; Liljencrantz and Olausson, 2014). In addition to this crucial observation, we provide the first experimental evidence that C-LTMRs contribute to evoked, affective pain after SCI. By utilizing a novel and feasible behavioral paradigm, in combination with optogenetic technology, we show that C-LTMR activation is sufficient for at-level neuropathic pain after SCI. Moreover, we report a complex interaction between C-LTMR stimulation and sympathetic activation, assessed through RRs. Like affective pain, RRs were elevated post-injury. Together, these findings bolster previous reports that sympathetic activity exerts influence over sensory systems in conditions of pain hypersensitivity. Counteractive ‘parasympathetic’ manipulations such as slowing respiration could provide a voluntary portal to autonomic nervous system control (Noble and Hochman, 2019) to reduce pain hypersensitivity, potentially by directly inhibiting C-LTMRs. For instance, operantly training rodents to slow their breathing decreases sensitivity to painful stimuli (Noble et al., 2017), and is one promising manipulation that may warrant further investigation. Finally, the methods and outcomes of this study provide a foundation for investigating the peripheral and central mechanisms of SCI-induced pain that could ultimately translate into new therapeutic options for pain control in humans.

## Acknowledgements

The authors would like to thank William N. Goolsby (Emory University) for technical help including constructing the CPA device, providing remote (EPIC) sensors for experimental use, and developing the LabVIEW program for real-time RR monitoring and data collection; and Shawn Hochman and Makalele Provost (Emory University) for undertaking electrophysiological recordings. The MMRRC at JAX is supported by the Office of Research Infrastructure Programs/Office of the Director (grant number OD010921) of the National Institutes of Health.

## References

Andrew, D. (2010). Quantitative characterization of low-threshold mechanoreceptor inputs to lamina I spinoparabrachial neurons in the rat. J Physiol 588(Pt 1), 117–124. doi: 10.1113/jphysiol.2009.181511.

Baastrup, C., and Finnerup, N.B. (2008). Pharmacological management of neuropathic pain following spinal cord injury. CNS Drugs 22(6), 455–475.

Bagdas, D., Muldoon, P.P., AlSharari, S., Carroll, F.I., Negus, S.S., and Damaj, M.I. (2016). Expression and pharmacological modulation of visceral pain-induced conditioned place aversion in mice. Neuropharmacology 102, 236–243. doi: 10.1016/j.neuropharm.2015.11.024.

Barasi, S., and Lynn, B. (1986). Effects of sympathetic stimulation on mechanoreceptive and nociceptive afferent units from the rabbit pinna. Brain Res 378(1), 21–27.

Basso, D.M., Fisher, L.C., Anderson, A.J., Jakeman, L.B., McTigue, D.M., and Popovich, P.G. (2006). Basso Mouse Scale for locomotion detects differences in recovery after spinal cord injury in five common mouse strains. J Neurotrauma 23(5), 635–659. doi: 10.1089/neu.2006.23.635.

Bedi, S.S., Yang, Q., Crook, R.J., Du, J., Wu, Z., Fishman, H.M., et al. (2010). Chronic spontaneous activity generated in the somata of primary nociceptors is associated with pain-related behavior after spinal cord injury. J Neurosci 30(44), 14870–14882. doi: 10.1523/JNEUROSCI.2428-10.2010.

Bessou, P., Burgess, P.R., Perl, E.R., and Taylor, C.B. (1971). Dynamic properties of mechanoreceptors with unmyelinated (C) fibers. J Neurophysiol 34(1), 116–131.

Chamessian, A., Matsuda, M., Young, M., Wang, M., Zhang, Z.J., Liu, D., et al. (2019). Is Optogenetic Activation of Vglut1-positive Abeta Low-Threshold Mechanoreceptors Sufficient to Induce Tactile Allodynia in Mice after Nerve Injury? J Neurosci. doi: 10.1523/JNEUROSCI.2064-18.2019.

Chaplan, S.R., Bach, F.W., Pogrel, J.W., Chung, J.M., and Yaksh, T.L. (1994). Quantitative assessment of tactile allodynia in the rat paw. J Neurosci Methods 53(1), 55–63.

Christensen, M.D., and Hulsebosch, C.E. (1997). Chronic central pain after spinal cord injury. J Neurotrauma 14(8), 517–537. doi: 10.1089/neu.1997.14.517.

Chung, K., Lee, B.H., Yoon, Y.W., and Chung, J.M. (1996). Sympathetic sprouting in the dorsal root ganglia of the injured peripheral nerve in a rat neuropathic pain model. J Comp Neurol 376(2), 241–252. doi: 10.1002/(SICI)1096-9861(19961209)376:2<241::AID-CNE6>3.0.CO;2-3.

Craig, A.D. (2010). Why a soft touch can hurt. J Physiol 588(Pt 1), 13. doi: 10.1113/jphysiol.2009.185116.

Crown, E.D., Ye, Z., Johnson, K.M., Xu, G.Y., McAdoo, D.J., and Hulsebosch, C.E. (2006). Increases in the activated forms of ERK 1/2, p38 MAPK, and CREB are correlated with the expression of at-level mechanical allodynia following spinal cord injury. Exp Neurol 199(2), 397–407. doi: 10.1016/j.expneurol.2006.01.003.

Culman, J., Ritter, S., Ohlendorf, C., Haass, M., Maser-Gluth, C., Spitznagel, H., et al. (1997). A new formalin test allowing simultaneous evaluation of cardiovascular and nociceptive responses. Can J Physiol Pharmacol 75(10-11), 1203–1211.

Delfini, M.C., Mantilleri, A., Gaillard, S., Hao, J., Reynders, A., Malapert, P., et al. (2013). TAFA4, a chemokine-like protein, modulates injury-induced mechanical and chemical pain hypersensitivity in mice. Cell Rep 5(2), 378–388. doi: 10.1016/j.celrep.2013.09.013.

Detloff, M.R., Smith, E.J., Quiros Molina, D., Ganzer, P.D., and Houle, J.D. (2014). Acute exercise prevents the development of neuropathic pain and the sprouting of non-peptidergic (GDNF-and artemin-responsive) c-fibers after spinal cord injury. Exp Neurol 255, 38–48. doi: 10.1016/j.expneurol.2014.02.013.

Felix, E.R., Cruz-Almeida, Y., and Widerstrom-Noga, E.G. (2007). Chronic pain after spinal cord injury: what characteristics make some pains more disturbing than others? J Rehabil Res Dev 44(5), 703–715.

Finnerup, N.B., Johannesen, I.L., Fuglsang-Frederiksen, A., Bach, F.W., and Jensen, T.S. (2003). Sensory function in spinal cord injury patients with and without central pain. Brain 126(Pt 1), 57–70.

Finnerup, N.B., Johannesen, I.L., Sindrup, S.H., Bach, F.W., and Jensen, T.S. (2001). Pain and dysesthesia in patients with spinal cord injury: A postal survey. Spinal Cord 39(5), 256–262. doi: 10.1038/sj.sc.3101161.

Gao, Y.J., and Ji, R.R. (2009). c-Fos and pERK, which is a better marker for neuronal activation and central sensitization after noxious stimulation and tissue injury? Open Pain J 2, 11–17. doi: 10.2174/1876386300902010011.

Garraway, S.M., Turtle, J.D., Huie, J.R., Lee, K.H., Hook, M.A., Woller, S.A., et al. (2011). Intermittent noxious stimulation following spinal cord contusion injury impairs locomotor recovery and reduces spinal brain-derived neurotrophic factor-tropomyosin-receptor kinase signaling in adult rats. Neuroscience 199, 86–102. doi: 10.1016/j.neuroscience.2011.10.007.

Garraway, S.M., Woller, S.A., Huie, J.R., Hartman, J.J., Hook, M.A., Miranda, R.C., et al. (2014). Peripheral noxious stimulation reduces withdrawal threshold to mechanical stimuli after spinal cord injury: role of tumor necrosis factor alpha and apoptosis. Pain 155(11), 2344–2359. doi: 10.1016/j.pain.2014.08.034.

Grant, J.A., and Rainville, P. (2009). Pain sensitivity and analgesic effects of mindful states in Zen meditators: a cross-sectional study. Psychosom Med 71(1), 106–114. doi: 10.1097/PSY.0b013e31818f52ee.

Habler, H.J., Janig, W., and Koltzenburg, M. (1987). Activation of unmyelinated afferents in chronically lesioned nerves by adrenaline and excitation of sympathetic efferents in the cat. Neurosci Lett 82(1), 35–40. doi: 10.1016/0304-3940(87)90167-4.

Hu, S., and Zhu, J. (1989). Sympathetic facilitation of sustained discharges of polymodal nociceptors. Pain 38(1), 85–90. doi: 10.1016/0304-3959(89)90077-8.

Hu, S.J., Yang, H.J., Jian, Z., Long, K.P., Duan, Y.B., Wan, Y.H., et al. (2000). Adrenergic sensitivity of neurons with non-periodic firing activity in rat injured dorsal root ganglion. Neuroscience 101(3), 689–698. doi: 10.1016/s0306-4522(00)00414-0.

Hummel, M., Lu, P., Cummons, T.A., and Whiteside, G.T. (2008). The persistence of a long-term negative affective state following the induction of either acute or chronic pain. Pain 140(3), 436–445. doi: 10.1016/j.pain.2008.09.020.

Iggo, A. (1960). Cutaneous mechanoreceptors with afferent C fibres. J Physiol 152, 337–353.

Jackson, A.B., and Groomes, T.E. (1994). Incidence of respiratory complications following spinal cord injury. Arch Phys Med Rehabil 75(3), 270–275.

Jänig, W., Levine, J.D., and Michaelis, M. (1996). “Chapter 10. Interactions of sympathetic and primary afferent neurons following nerve injury and tissue trauma,” in Progress in Brain Research, eds. L.K. Takao Kumazawa & M. Kazue. Elsevier), 161–184.

Kambrun, C., Roca-Lapirot, O., Salio, C., Landry, M., Moqrich, A., and Le Feuvre, Y. (2018). TAFA4 Reverses Mechanical Allodynia through Activation of GABAergic Transmission and Microglial Process Retraction. Cell Reports 22(11), 2886–2897. doi: 10.1016/j.celrep.2018.02.068.

Kinnman, E., and Levine, J.D. (1995a). Involvement of the sympathetic postganglionic neuron in capsaicin-induced secondary hyperalgesia in the rat. Neuroscience 65(1), 283–291.

Kinnman, E., and Levine, J.D. (1995b). Sensory and sympathetic contributions to nerve injury-induced sensory abnormalities in the rat. Neuroscience 64(3), 751–767.

Li, L., Rutlin, M., Abraira, V.E., Cassidy, C., Kus, L., Gong, S., et al. (2011). The functional organization of cutaneous low-threshold mechanosensory neurons. Cell 147(7), 1615–1627. doi: 10.1016/j.cell.2011.11.027.

Liljencrantz, J., Bjornsdotter, M., Morrison, I., Bergstrand, S., Ceko, M., Seminowicz, D.A., et al. (2013). Altered C-tactile processing in human dynamic tactile allodynia. Pain 154(2), 227–234. doi: 10.1016/j.pain.2012.10.024.

Liljencrantz, J., and Olausson, H. (2014). Tactile C fibers and their contributions to pleasant sensations and to tactile allodynia. Frontiers in Behavioral Neuroscience 8. doi: ARTN 37 10.3389/fnbeh.2014.00037.

Liljencrantz, J., Strigo, I., Ellingsen, D.M., Kramer, H.H., Lundblad, L.C., Nagi, S.S., et al. (2017). Slow brushing reduces heat pain in humans. Eur J Pain 21(7), 1173–1185. doi: 10.1002/ejp.1018.

Loggia, M.L., Juneau, M., and Bushnell, M.C. (2011). Autonomic responses to heat pain: Heart rate, skin conductance, and their relation to verbal ratings and stimulus intensity. Pain 152(3), 592–598. doi: 10.1016/j.pain.2010.11.032.

Loken, L.S., Wessberg, J., Morrison, I., McGlone, F., and Olausson, H. (2009). Coding of pleasant touch by unmyelinated afferents in humans. Nature Neuroscience 12(5), 547–548. doi: 10.1038/nn.2312.

Lou, S., Duan, B., Vong, L., Lowell, B.B., and Ma, Q. (2013). Runx1 controls terminal morphology and mechanosensitivity of VGLUT3-expressing C-mechanoreceptors. J Neurosci 33(3), 870–882. doi: 10.1523/JNEUROSCI.3942-12.2013.

Mahns, D.A., and Nagi, S.S. (2013). An investigation into the peripheral substrates involved in the tactile modulation of cutaneous pain with emphasis on the C-tactile fibres. Exp Brain Res 227(4), 457–465. doi: 10.1007/s00221-013-3521-5.

Martin, K.K., Noble, D.J., Parvin, S., Jang, K., and Garraway, S.M. (2022). Pharmacogenetic inhibition of TrkB signaling in adult mice attenuates mechanical hypersensitivity and improves locomotor function after spinal cord injury. Front Cell Neurosci 16, 987236. doi: 10.3389/fncel.2022.987236.

Martin, K.K., Parvin, S., and Garraway, S.M. (2019). Peripheral Inflammation Accelerates the Onset of Mechanical Hypersensitivity after Spinal Cord Injury and Engages Tumor Necrosis Factor alpha Signaling Mechanisms. J Neurotrauma 36(12), 2000–2010. doi: 10.1089/neu.2018.5953.

Nagi, S.S., and Mahns, D.A. (2013). C-tactile fibers contribute to cutaneous allodynia after eccentric exercise. J Pain 14(5), 538–548. doi: 10.1016/j.jpain.2013.01.009.

Nagi, S.S., Rubin, T.K., Chelvanayagam, D.K., Macefield, V.G., and Mahns, D.A. (2011). Allodynia mediated by C-tactile afferents in human hairy skin. J Physiol 589(Pt 16), 4065–4075. doi: 10.1113/jphysiol.2011.211326.

Neumann, S., Doubell, T.P., Leslie, T., and Woolf, C.J. (1996). Inflammatory pain hypersensitivity mediated by phenotypic switch in myelinated primary sensory neurons. Nature 384(6607), 360–364. doi: 10.1038/384360a0.

Noble, D.J., Goolsby, W.N., Garraway, S.M., Martin, K.K., and Hochman, S. (2017). Slow Breathing Can Be Operantly Conditioned in the Rat and May Reduce Sensitivity to Experimental Stressors. Frontiers in Physiology 8. doi: ARTN 854 10.3389/fphys.2017.00854.

Noble, D.J., and Hochman, S. (2019). Hypothesis: Pulmonary Afferent Activity Patterns During Slow, Deep Breathing Contribute to the Neural Induction of Physiological Relaxation. Front Physiol 10, 1176. doi: 10.3389/fphys.2019.01176.

Noble, D.J., Martin, K.K., Parvin, S., and Garraway, S.M. (2019). Spontaneous and Stimulus-Evoked Respiratory Rate Elevation Corresponds to Development of Allodynia in Spinal Cord-Injured Rats. J Neurotrauma. doi: 10.1089/neu.2018.5936.

Oatway, M.A., Chen, Y., and Weaver, L.C. (2004). The 5-HT3 receptor facilitates at-level mechanical allodynia following spinal cord injury. Pain 110(1-2), 259–268. doi: 10.1016/j.pain.2004.03.040.

Olausson, H., Lamarre, Y., Backlund, H., Morin, C., Wallin, B.G., Starck, G., et al. (2002). Unmyelinated tactile afferents signal touch and project to insular cortex. Nat Neurosci 5(9), 900–904. doi: 10.1038/nn896.

Parvin, S., Williams, C.R., Jarrett, S.A., and Garraway, S.M. (2021). Spinal Cord Injury Increases Pro-inflammatory Cytokine Expression in Kidney at Acute and Sub-chronic Stages. Inflammation 44(6), 2346–2361. doi: 10.1007/s10753-021-01507-x.

Pertens, E., Urschel-Gysbers, B.A., Holmes, M., Pal, R., Foerster, A., Kril, Y., et al. (1999). Intraspinal and behavioral consequences of nerve growth factor-induced nociceptive sprouting and nerve growth factor-induced hyperalgesia compared in adult rats. J Comp Neurol 410(1), 73–89.

Pinheiro Fde, V., Villarinho, J.G., Silva, C.R., Oliveira, S.M., Pinheiro Kde, V., Petri, D., et al. (2015). The involvement of the TRPA1 receptor in a mouse model of sympathetically maintained neuropathic pain. Eur J Pharmacol 747, 105–113. doi: 10.1016/j.ejphar.2014.11.039.

Provost, M.A. (2019). The effects of adrenergic, serotonergic, and cholinergic modulation on hairy skin low threshold mechanoreceptors. Master of Science, Emory University.

Ramer, M.S., and Bisby, M.A. (1997). Rapid sprouting of sympathetic axons in dorsal root ganglia of rats with a chronic constriction injury. Pain 70(2–3), 237–244. doi: http://dx.doi.org/10.1016/S0304-3959(97)03331-9.

Ramer, M.S., Thompson, S.W.N., and McMahon, S.B. (1999). Causes and consequences of sympathetic basket formation in dorsal root ganglia. Pain 82, Supplement 1(0), S111–S120. doi: http://dx.doi.org/10.1016/S0304-3959(99)00144-X.

Refsgaard, L.K., Hoffmann-Petersen, J., Sahlholt, M., Pickering, D.S., and Andreasen, J.T. (2016). Modelling affective pain in mice: Effects of inflammatory hypersensitivity on place escape/avoidance behaviour, anxiety and hedonic state. J Neurosci Methods 262, 85–92. doi: 10.1016/j.jneumeth.2016.01.019.

Roberts, W.J., and Elardo, S.M. (1985). Sympathetic activation of unmyelinated mechanoreceptors in cat skin. Brain Res 339(1), 123–125.

Roberts, W.J., and Levitt, G.R. (1982). Histochemical evidence for sympathetic innervation of hair receptor afferents in cat skin. J Comp Neurol 210(2), 204–209. doi: 10.1002/cne.902100211.

Ruocco, I., Cuello, A.C., and Ribeiro-Da-Silva, A. (2000). Peripheral nerve injury leads to the establishment of a novel pattern of sympathetic fibre innervation in the rat skin. The Journal of Comparative Neurology 422(2), 287–296. doi: 10.1002/(SICI)1096-9861(20000626)422:2<287::AID-CNE9>3.0.CO;2-E.

Santuzzi, C.H., Neto Hde, A., Pires, J.G., Goncalves, W.L., Gouvea, S.A., and Abreu, G.R. (2013). High-frequency transcutaneous electrical nerve stimulation reduces pain and cardio-respiratory parameters in an animal model of acute pain: participation of peripheral serotonin. Physiother Theory Pract 29(8), 630–638. doi: 10.3109/09593985.2013.774451.

Sato, J., and Perl, E.R. (1991). Adrenergic excitation of cutaneous pain receptors induced by peripheral nerve injury. Science 251(5001), 1608–1610. doi: 10.1126/science.2011742.

Sato, J., Suzuki, S., Iseki, T., and Kumazawa, T. (1993). Adrenergic excitation of cutaneous nociceptors in chronically inflamed rats. Neurosci Lett 164(1-2), 225–228. doi: 10.1016/0304-3940(93)90897-t.

Seal, R.P., Wang, X., Guan, Y., Raja, S.N., Woodbury, C.J., Basbaum, A.I., et al. (2009). Injury-induced mechanical hypersensitivity requires C-low threshold mechanoreceptors. Nature 462(7273), 651–655. doi: 10.1038/nature08505.

Shinder, V., Govrin-Lippmann, R., Cohen, S., Belenky, M., Ilin, P., Fried, K., et al. (1999). Structural basis of sympathetic-sensory coupling in rat and human dorsal root ganglia following peripheral nerve injury. J Neurocytol 28(9), 743–761. doi: 10.1023/a:1007090105840.

Siddall, P.J., and Loeser, J.D. (2001). Pain following spinal cord injury. Spinal Cord 39(2), 63–73.

Siddall, P.J., McClelland, J.M., Rutkowski, S.B., and Cousins, M.J. (2003). A longitudinal study of the prevalence and characteristics of pain in the first 5 years following spinal cord injury. Pain 103(3), 249–257.

Tanimoto, K., Takebayashi, T., Kobayashi, T., Tohse, N., and Yamashita, T. (2011). Does norepinephrine influence pain behavior mediated by dorsal root ganglia?: a pilot study. Clin Orthop Relat Res 469(9), 2568–2576. doi: 10.1007/s11999-011-1798-x.

Teng, Y.D., Bingaman, M., Taveira-DaSilva, A.M., Pace, P.P., Gillis, R.A., and Wrathall, J.R. (2003). Serotonin 1A receptor agonists reverse respiratory abnormalities in spinal cord-injured rats. J Neurosci 23(10), 4182–4189.

Teng, Y.D., Mocchetti, I., Taveira-DaSilva, A.M., Gillis, R.A., and Wrathall, J.R. (1999). Basic fibroblast growth factor increases long-term survival of spinal motor neurons and improves respiratory function after experimental spinal cord injury. J Neurosci 19(16), 7037–7047.

Vrontou, S., Wong, A.M., Rau, K.K., Koerber, H.R., and Anderson, D.J. (2013). Genetic identification of C fibres that detect massage-like stroking of hairy skin in vivo. Nature 493(7434), 669–+. doi: 10.1038/nature11810.

Weaver, L.C., Verghese, P., Bruce, J.C., Fehlings, M.G., Krenz, N.R., and Marsh, D.R. (2001). Autonomic dysreflexia and primary afferent sprouting after clip-compression injury of the rat spinal cord. J Neurotrauma 18(10), 1107–1119. doi: 10.1089/08977150152693782.

Wolf, S., and Hardy, J.D. (1941). Studies on Pain. Observations on Pain Due to Local Cooling and on Factors Involved in the “Cold Pressor” Effect. J Clin Invest 20(5), 521–533. doi: 10.1172/JCI101245.

Wu, Z., Li, L., Xie, F., Du, J., Zuo, Y., Frost, J.A., et al. (2017). Activation of KCNQ Channels Suppresses Spontaneous Activity in Dorsal Root Ganglion Neurons and Reduces Chronic Pain after Spinal Cord Injury. J Neurotrauma 34(6), 1260–1270. doi: 10.1089/neu.2016.4789.

Xu, Q., Garraway, S.M., Weyerbacher, A.R., Shin, S.J., and Inturrisi, C.E. (2008). Activation of the neuronal extracellular signal-regulated kinase 2 in the spinal cord dorsal horn is required for complete Freund’s adjuvant-induced pain hypersensitivity. J Neurosci 28(52), 14087–14096. doi: 10.1523/JNEUROSCI.2406-08.2008.

Yang, Q., Wu, Z.Z., Hadden, J.K., Odem, M.A., Zuo, Y., Crook, R.J., et al. (2014). Persistent Pain after Spinal Cord Injury Is Maintained by Primary Afferent Activity. Journal of Neuroscience 34(32), 10765–10769. doi: 10.1523/Jneurosci.5316-13.2014.

Yezierski, R.P., Yu, C.G., Mantyh, P.W., Vierck, C.J., and Lappi, D.A. (2004). Spinal neurons involved in the generation of at-level pain following spinal injury in the rat. Neurosci Lett 361(1-3), 232–236. doi: 10.1016/j.neulet.2003.12.035.

Zhang, J.-M., Li, H., and Munir, M.A. (2004). Decreasing sympathetic sprouting in pathologic sensory ganglia: a new mechanism for treating neuropathic pain using lidocaine. Pain 109(1–2), 143–149. doi: http://dx.doi.org/10.1016/j.pain.2004.01.033.

Zheng, Q., Xie, W., Luckemeyer, D.D., Lay, M., Wang, X.W., Dong, X., et al. (2022). Synchronized cluster firing, a distinct form of sensory neuron activation, drives spontaneous pain. Neuron 110(2), 209–220 e206. doi: 10.1016/j.neuron.2021.10.019.

Zimmerman, A., Bai, L., and Ginty, D.D. (2014). The gentle touch receptors of mammalian skin. Science 346(6212), 950–954. doi: 10.1126/science.1254229.

